# Generation of a single-cell B cell atlas of antibody repertoires and transcriptomes to identify signatures associated with antigen-specificity

**DOI:** 10.1101/2021.11.09.467876

**Authors:** Andreas Agrafiotis, Daniel Neumeier, Kai-Lin Hong, Tasnia Chowdhury, Roy Ehling, Raphael Kuhn, Ioana Sandu, Victor Kreiner, Tudor-Stefan Cotet, Daria Laslo, Stine Anzböck, Dale Starkie, Daniel J. Lightwood, Annette Oxenius, Sai T. Reddy, Alexander Yermanos

## Abstract

Murine models of immunization have played a major role in discovering antibody candidates against therapeutic targets. It nevertheless remains time-consuming and expensive to identify antibodies with diverse binding modalities against druggable candidate molecules. Although new genomics-based pipelines have potential to augment antibody discovery, these methods remain in their infancy due to an incomplete understanding of the selection process that governs B cell clonal selection, expansion and antigen specificity. Furthermore, it remains unknown how factors such as aging and reduction of tolerance influence B cell selection in murine models of immunization. Here we perform single-cell sequencing of antibody repertoires and transcriptomes of B cells following immunizations with a model therapeutic antigen target (human Tumor necrosis factor receptor 2, TNFR2). We determine the relationship between antibody repertoires, gene expression signatures and antigen specificity across 100,000 B cells. Recombinant expression and characterization of 227 monoclonal antibodies revealed the existence of clonally expanded and class-switched antigen-specific B cells that were more frequent in young mice. Although integrating multiple repertoire features such as germline gene usage, somatic hypermutation, and transcriptional signatures failed to distinguish antigen-specific from non-specific B cells, other features such as IgG-subtype and sequence composition correlated with antigen-specificity. This work provides a single-cell resource for B cells relating antibody repertoires, transcriptomes and antigen specificity.

## Introduction

The importance of antibodies in drug development has promoted interest in developing more rapid and efficient methods for monoclonal antibody discovery (Huggett 2018; Weiner 2015). Traditionally antibody discovery has relied heavily on experimental screening approaches such as hybridomas, phage or yeast display and more recently B cell cloning (Hoogenboom 2005; Wilson and Andrews 2012); however the emergence of genomics methods and in particular deep sequencing and bioinformatics are also contributing to antibody discovery (C. Parola, Neumeier, and Reddy 2018). For example, deep sequencing of antibody repertoires from immunized mice has been used to identify clonally expanded plasma cells that are associated with antigen specificity (Reddy et al. 2010; Wang et al. 2015). Additionally, the emergence of single-cell sequencing (scSeq) technologies has made it possible to identify endogenously paired variable heavy (VH) and variable light chain (VL) regions from single B cells at high-throughput, which can be used to subsequently reconstruct antibodies for experimental testing <u>(DeKosky et al. 2013; DeKosky et al. 2015)</u>. For example, our group recently performed scSeq of plasma cells following immunization or viral infection in mice, and subsequent expression and screening of antibodies from highly expanded clones was shown to correlate with antigen-specificity (Neumeier et al. 2022; Neumeier, Yermanos, et al. 2021). A similar approach was applied to plasma cells isolated from convalescent Covid-19 individuals and led to the discovery of a panel of antibodies, including potent neutralizing antibodies against SARS-CoV-2 (Ehling et al. 2022). An innovative approach based on B cell selection with DNA-barcoded antigens followed by scSeq (LIBRA-seq) has been effective for the rapid isolation of antibodies against multiple antigens (Setliff et al. 2019; Shiakolas et al. 2022).

Further development of scSeq methods has made it possible to obtain both transcriptome and antibody (or B cell receptor, BCR) repertoire information from single B cells at high-throughput (Croote et al. 2018; Horns, Dekker, and Quake 2020; Saikia et al. 2019; Singh et al. 2019), thereby enabling an integrated analysis of B cell transcriptional phenotypes within the context of antibody clonal populations (Yermanos, Agrafiotis, et al. 2021). This integrated scSeq approach was recently used to identify a large and diverse population of antigen-specific B cells from convalescent Covid-19 individuals, leading to the discovery of highly potent neutralizing antibodies (Cao et al. 2020). Although this work demonstrates the promise of integrated scSeq for viral antigens, it remains unknown how antibody repertoire and transcriptome features can be used to identify antigen specific B cells, especially within the context of challenging therapeutic target antigens, such as self-antigens that are often targets for autoimmune disease or cancer (Fischer, Kontermann, and Pfizenmaier 2020;Medler and Wajant 2019). The discovery of antigen specific B cells to antigens with high homology to self proteins may be potentially enhanced through immunization in aged individuals, as previous studies have demonstrated that the redistribution of B cell subsets that occurs during aging may favor the formation of antibody responses towards self-antigens (Frasca et al. 2017; Colonna-Romano et al. 2009; Hao et al. 2011). Furthermore, it remains unknown whether the potential loss of negative selection occurring with immunosenescence can improve the ability to identify diverse populations of antigen-specific B cells (Rubtsov et al. 2011; Watad et al. 2017; Dunn-Walters 2015).

Here, we perform integrated scSeq of antibody repertoires and transcriptomes of B cells isolated from mice immunized with a model therapeutic protein antigen (human TNFR2) followed by antibody-antigen specificity profiling at high-throughput. We generate a single-cell atlas of ~100,000 B cells and experimentally screen 227 antibodies, thereby uncovering signatures associated with antigen-specific B cells. Importantly, only in young mice were clonally expanded B cells found to be strongly associated with antigen-specific antibodies, with very few observed antigen-specific B cells discovered in the older cohort. In addition to supporting the discovery of a diverse panel of antigen-specific B cells against a therapeutically relevant target, scSeq can also relate functional properties of antibodies to repertoire and transcriptional features and gain greater insight on B cell selection.

## Results

### Clonal expansion is detected in young and old mice following immunization

To profile B cell selection following serial protein immunizations, we immunized a cohort of 3-month-old (3m) male C57BL6 mice (n=5) with five successive injections of 10 μg of the extracellular domain of human TNFR2 mixed with 20 μg of the adjuvant monophosphoryl lipid A (MPLA). Our previous findings demonstrated that immunization of young mice with an immunogenic protein antigen (ovalbumin) results in highly expanded plasma cells within the bone marrow (Neumeier, Yermanos, et al. 2021). For this study, we selected an antigen that has potentially reduced immunogenicity given its high sequence similarity (~61.3%) to the murine homologue of TNFR2. As aging may be linked to a more antigen experienced immune repertoire (encounter of higher number of antigens throughout life) together with an increased production of autoantibodies (Ratliff et al. 2013; Elkon and Casali 2008), we hypothesized that immunization with TNFR2 in aged mice could potentially trigger stronger memory B cell responses against self antigens with high sequence similarity, such as human TNFR2. Furthermore, it has been observed that antibody forming B cells in aged mice exhibit higher absolute numbers in certain organs following cognate antigen interactions (Han et al. 2003). Hence, we additionally performed the same immunization schemes with additional cohorts of 12-month-old (12m) (n=3) and 18-month-old (18m) (n=3) male C57BL6 mice. With the exception of one mouse in the young cohort, all mice exhibited high antibody titers against TNFR2 (Figure S1).

We isolated bone marrow plasma cells (BM PCs) (CD138^hi^, TACI^hi^) and spleen B cells (CD19^hi^, IgM^low^, IgD^low^) by fluorescence activated cell sorting (FACS) and performed single-cell antibody repertoire and transcriptome sequencing using the 10x genomics 5’ immune (VDJ) profiling pipeline (Figure 1A). Following library preparation, deep sequencing and alignment to reference VDJ genes, we recovered a total of ~65,000 cells containing exactly one heavy chain and one light chain, with an average of 3,200 cells per mouse. The majority of cells were of the IgM isotype for both spleen and bone marrow (Figure 1B). Cells were grouped together based on sharing identical heavy and light chain complementarity determining region 3 (CDRH3+CDRL3) amino acid sequences (hereafter referred to as clone), resulting in a few hundreds of unique clones for each mouse or organ repertoire. Further investigation revealed that the majority of the BM PC repertoires demonstrated extensive clonal expansion, with approximately 75% of clones being expanded (clones supported by more than one cell) (Figures 1D, S2). On the other hand, we observed that the majority of splenic B cell clones were not clonally expanded, as they possessed only a single cell barcode, which was comparable across all three age groups (Figures 1D, S2). While the majority of expanded clones for BM PCs were IgM, spleen repertoires had an extensive presence of IgG subtypes (Figures 1E, S3, S4). Nevertheless, clonally expanded cells of IgM, IgG and IgA isotypes were still present in both organs (Figures S5, S6, S7). Previous single-cell antibody repertoire sequencing studies suggest a strong correlation between class-switched clones and the number of amino acid sequence variants existing within an individual clone (Neumeier, Pedrioli, et al. 2021; Neumeier, Yermanos, et al. 2021). Here, we observed that the number of cells for the most expanded IgG and IgA clones demonstrated a minor correlation with the number of amino acid variants, whereas some IgM clones contained a considerable number of amino acid variants (Figure 1F). Finally, we did not observe any differences in clonal expansion, isotype distribution or the number of amino acid variants between the different age groups.

**Figure 1.**
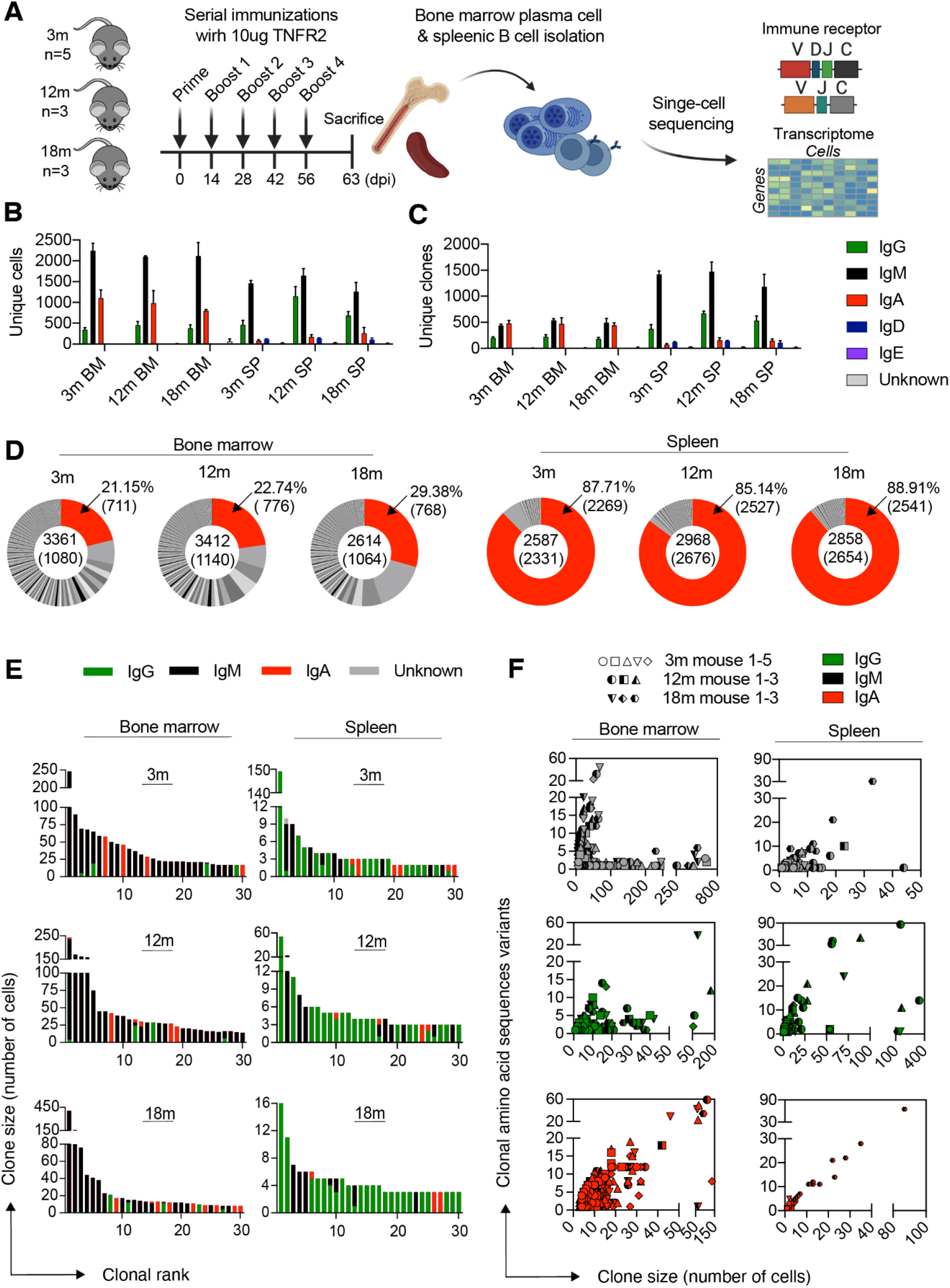
Single-cell antibody repertoire sequencing reveals clonal expansion and class-switching in bone marrow plasma cells and splenic B cells following immunization. A. Experimental overview of immunization with antigen (TNFR2), bone marrow plasma cell (BM PC) and splenic B cell isolation, single-cell antibody repertoire sequencing. B. Mean number of cells per isotype for each immunized cohort. Only cells containing exactly one variable heavy (VH) and variable light (VL) chain were retained in the analysis. Colors correspond to isotype. C. Mean number of clones per isotype for each immunized cohort. Clones were determined by grouping B cells containing identical CDRH3+CDRL3 amino acid sequences. The isotype was determined as the isotype corresponding to the majority of cells within one clone. Color corresponds to isotype. D. Distribution of clonal expansion. Each section corresponds to a unique clone and the size corresponds to the fraction of cells relative to the total repertoire. Red color highlights the fraction of clones containing one cell. Numbers in the center indicate the total number of cells and clones (in parenthesis). Numbers on the right indicate the percentage and total number (in parenthesis) of unexpanded clones. One representative mouse per age and organ cohort is shown. E. Clonal frequency for the 30 most expanded clones in each repertoire. One representative mouse per age and organ cohort is shown. Clones were determined by grouping B cells containing identical CDRH3+CDRL3 amino acid sequences. Color corresponds to isotype. F. Relationship between the number of unique amino acid variants and the number of cell barcodes for the 30 most expanded clones separated by isotype majority.

### Single-cell sequencing reveals organ and isotype transcriptional heterogeneity

Next, we leveraged the capabilities of scSeq to integrate transcriptome information with antibody repertoire features. To this end, we performed unsupervised clustering and uniform manifold approximation projection (UMAP) on the combined data set (all cells across all mice and cohorts), which gave rise to 14 distinct cell clusters based on global gene expression (Figures 2A, S9A, S10). Examination of the transcriptional space clearly distinguishes between clusters populated principally by cells derived from either bone marrow or spleen, or clusters with cells originating from both organs at similar ratios (clusters 8, 9 and 12) (Figure 2A, S9B); a high degree of reproducibility was observed across samples (Figure S9C). Differential gene expression analysis was used to define specific cell populations (Figures 2B, S11) (Mathew et al. 2021). Not surprisingly, this demonstrated that cell clusters associated with bone marrow were characterized almost completely by plasma cell markers, which was in contrast to the splenic transcriptional space where multiple distinct B cell phenotypes were present (Figure 2A). Overlaying isotype information onto the UMAP revealed separation of IgG-, IgM-, IgA- and IgD-expressing cells (Figure S12). Differential gene expression analysis revealed exclusive genes defining IgG (*ApoE, Gimap4*), IgA (*Ccr10, Glpr1*), IgD (*Sell, Fcer2a*) and IgM (*Ggh, Slc3a2*) isotypes, some of which are consistent with previous findings (Neumeier, Pedrioli, et al. 2021) (Figure 2C).

**Figure 2.**
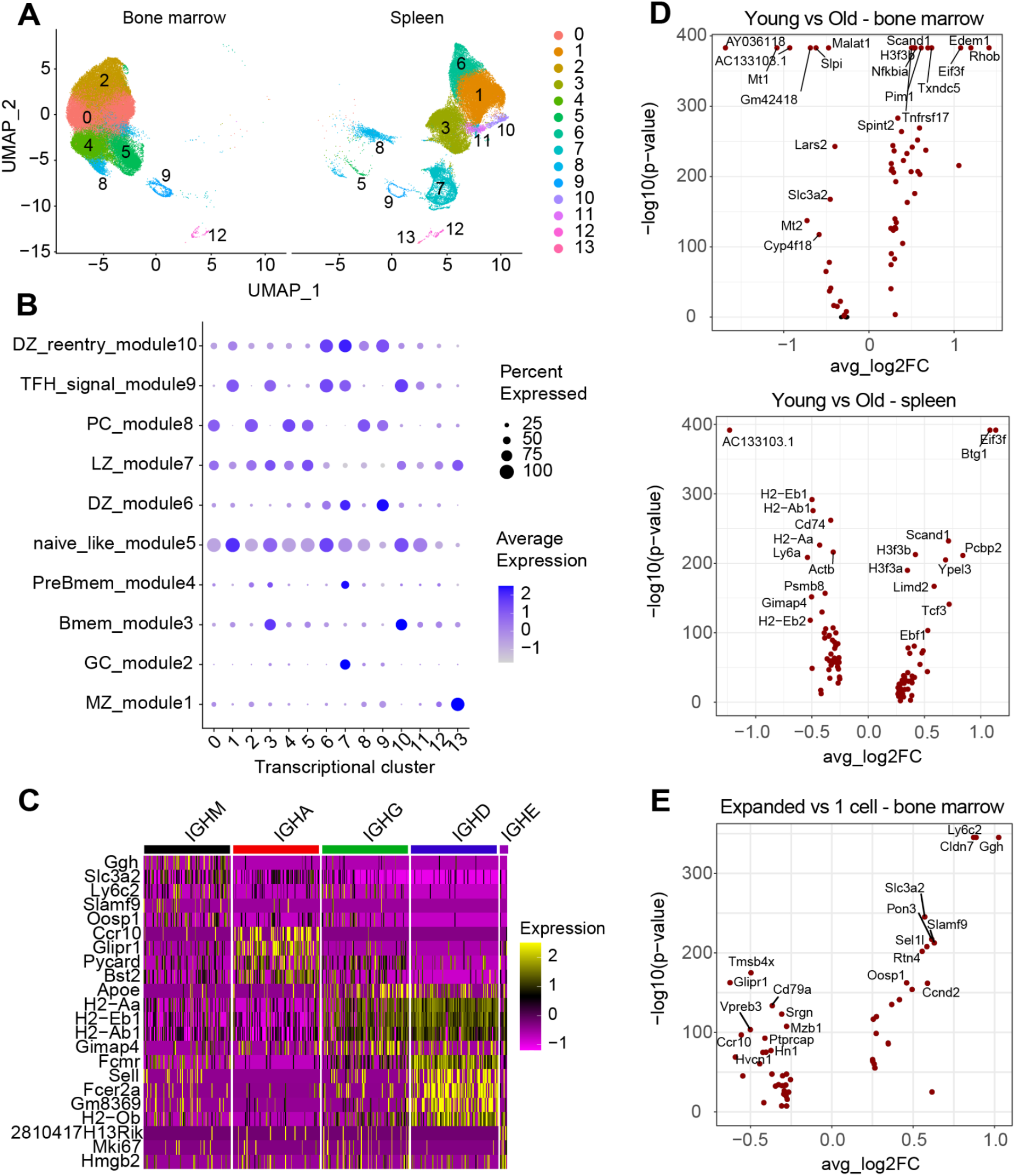
Single-cell transcriptome sequencing of B cells reveals organ, expansion, age and isotype transcriptional heterogeneity. A. Uniform manifold approximation projection (UMAP) based on total gene expression of all repertoires (split by organ) following immunization. Each point corresponds to a cell and color corresponds to the transcriptional cluster. B. Dottile plot showing B cell subset assignment across all clusters based on expression of genes defining B cell phenotypes. The intensity of each dot corresponds to the average expression of all cells within a given cluster and the size corresponds to the percentage of cells with detectable gene expression. C. Heatmap of differentially expressed genes between isotypes. D. Differential gene expression between young and old for bone marrow (top volcano plot) and spleen (bottom volcano plot). Points in red indicate significantly differentially expressed genes (p-adj < 0.01). E. Differential gene expression between expanded and single-cell clones in the bone marrow Points in red indicate significantly differentially expressed genes (p-adj < 0.01).

We next determined if there are age-associated transcriptional changes between young and old mice. We restricted our analysis to the 3m and 18m cohorts and quantified differential gene expression separately for each organ (Figure 2D). Interestingly, we observed for both bone marrow and spleen a substantial downregulation in young mice of *AC133103.1*, an uncharacterised gene marker, and substantial upregulation of *Eif3f* which has been implicated in protein translation and cell growth with its expression significantly decreased in many human cancers (Marchione, Leibovitch, and Lenormand 2013; Shi et al. 2006) (Figure 2D). We determined if expanded B cell clones were transcriptionally distinct compared to non-expanded clones (clones supported by only one single cell). Due to the high heterogeneity of the splenic B cell population, we focused our analysis on BM PCs (Figure 2E). Identification of cluster defining genes in the expanded BM PC pool was not obvious, however, we did observe a downregulation of *Vpreb3*, which is implicated in B cell maturation (Rodig et al. 2010;Rosnet et al. 2004) (Figure 2E).

### Clonal expansion correlates with antigen specificity in young but not old mice

We next determined the extent to which clonal expansion is correlated with antigen-specific binding. We focused on the most expanded clones that possessed cells mostly with an IgG subtype, as these have been previously shown to be preferentially associated with antigen specificity (Neumeier, Yermanos, et al. 2021). We used a previously established system for rapid antibody cloning and expression in mammalian cells by CRISPR-Cas9 genome editing (Cristina Parola et al. 2019; Pogson et al. 2016). The resulting antibodies, representing 204 unique IgG clones, were then screened by ELISA for binding to the TNFR2 antigen (Figure 3A, Table S1). Unexpectedly, we observed that in young mice, there was a considerably higher fraction of antigen-specific clones compared to older mice (Figures 3B, 3C, Table S1). This was most apparent in BM PC repertoires, where approximately 31.3% of clones tested (31 out of 99) from young mice were antigen-specific, in contrast to only 7.1% (4 out of 56) in older mice (Figure 3B). In the spleen we also detected 21.4% (6 out of 28) and 14.2% (3 out of 21) antigen-specific clones for young and old mice, respectively (Figure 3C). Furthermore, independent of age and organ, we observed that a higher degree of clonal expansion was not correlated with antigen specificity, as antigen-specific and non-specific clones were evenly distributed throughout the most expanded clones of each repertoire (Figures 3B, 3C).

**Figure 3.**
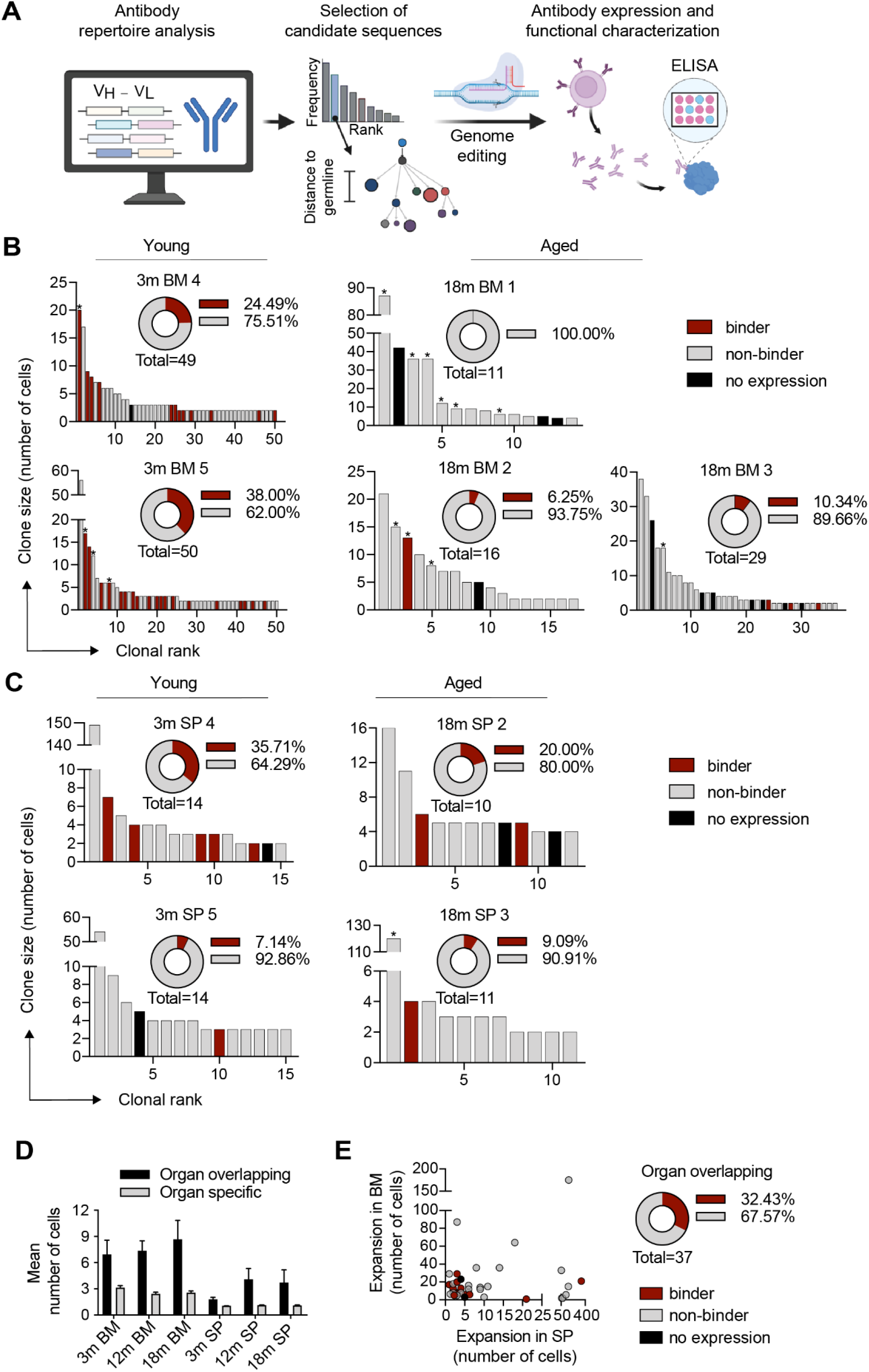
Antibody expression and antigen-specific screening. A. Workflow for interrogating the antibody specificity from immunized mice. Antibody repertoire analysis is performed to identify expanded class-switched clonal lineages, which are then used as templates for genome editing, through homology-directed recombination, and expression of antibody. Supernatant is used to determine reactivity of antibodies against immunizing molecule. B. & C. Clonal rank plot of most expanded IgG clones indicating TNFR2-specific clones in young and old bone marrow (B) and splenic (C) repertoires. Clones were determined by grouping B cells based on shared CDRH3+CDRL3 sequences (100% a.a. identity). Only cells containing exactly one variable heavy (VH) and variable light (VL) chain were considered. For each clone, the antibody variant (combined VH+VL nucleotide sequence) supported by the most unique cell barcodes was selected to be expressed. Sequences that also belong to overlapping clones are labeled with a star (*). D. Mean clonal expansion between organ-overlapping and organ-specific clones. E. Relationship between clonal expansion (number of cells) in the bone marrow (BM) (y-axis) and in the spleen (SP) (x axis) for clones present in both organs within the same mouse; a subset of clones were expressed as recombinant antibodies and binding to antigen (TNFR2) was determined. “No expression” corresponds to clones with antibody sequences that could not be expressed following immunogenic engineering of hybridoma cells.

Having discovered TNFR2-specific antibody sequences among clonally expanded cells, we next determined whether clones present in multiple organs of the same mouse would exhibit antigen-specificity. These organ-overlapping clones showed higher levels of clonal expansion when compared to clones observed in only one organ (Figure 3D). We expressed and determined the antigen specificity of several of these organ-overlapping clones, some of which coincided with previously expressed expanded clones (marked with a star(*) in Figures 3B and 3C). This revealed that approximately 33% (12/37) of organ-overlapping clones were also antigen-specific. (Figure 3E, Table S1). Furthermore, antigen specificity and clonal expansion were not correlated across organ-overlapping clones (Figure 2E). In contrast to the correlation observed between age and antigen-specificity in the clonally expanded cells, we did not observe any age associated biases to antigen specificity within the organ-overlapping antibody pool.

### Investigating immune repertoire and transcriptome features of antigen-specific BM PCs

We next determined whether repertoire or transcriptome features could differentiate between recently activated TNFR2-specific clones and non-specific ones. We focused our computational analysis on the BM PCs due to the higher number of experimentally validated antibodies with specificity to TNFR2. We initially quantified the mean number of somatic hypermutations (SHMs) per clone in the full-length VH and VL regions, which revealed that, on average, the TNFR2-specific fraction of clones exhibited lower levels of SHM (Figure 4A). Next, we visualized the distribution of IgG subtypes (IgG1, IgG1B, IgG2C, IgG3) which showed a higher percentage of the IgG1 subtype among the antigen-specific fraction of clones (Figures 4B), an observation that was also reflected on the cellular level (Figure S13A). We next sought to relate TNFR2 specificity to germline gene usage. Visualizing the distribution of heavy chain (HC) and light chain (LC) V gene usage across TNFR2-specific and non-specific sequences did not suggest any TNFR2-associated biases (Figure S13B). We further determined if certain VH-VL germline combinations were enriched in the antigen-specific or non-specific fraction. Circos plots of the experimentally verified antigen-specific and non-specific clones did not show enrichment for certain V gene combinations in either of the two groups (Figure 4C). The same observation was made when looking at other germline features, such as the J gene usage (Figure S13C). To investigate whether sequence convergence could be detected within the binding fraction of TNFR2-specific clones, we initially visualized the amino acid sequences for the most frequent CDRH3 and CDRL3 lengths. This initially revealed little indication of amino acid or biochemical bias between antigen-specific and non-specific sequences within the CDR3 region (Figures 4D, S13D). Next, we constructed sequence similarity networks based on the edit distance of CDRs (Miho et al. 2018), which demonstrated that antigen-specific clones demonstrated clustering across a range of edit distances (Figures 4E, S13E). Some of these antigen-specific clusters contained clones utilizing different germline genes despite similar CDR3s, potentially suggesting convergent sequence motifs (Figures 4F, S13F).

**Figure 4.**
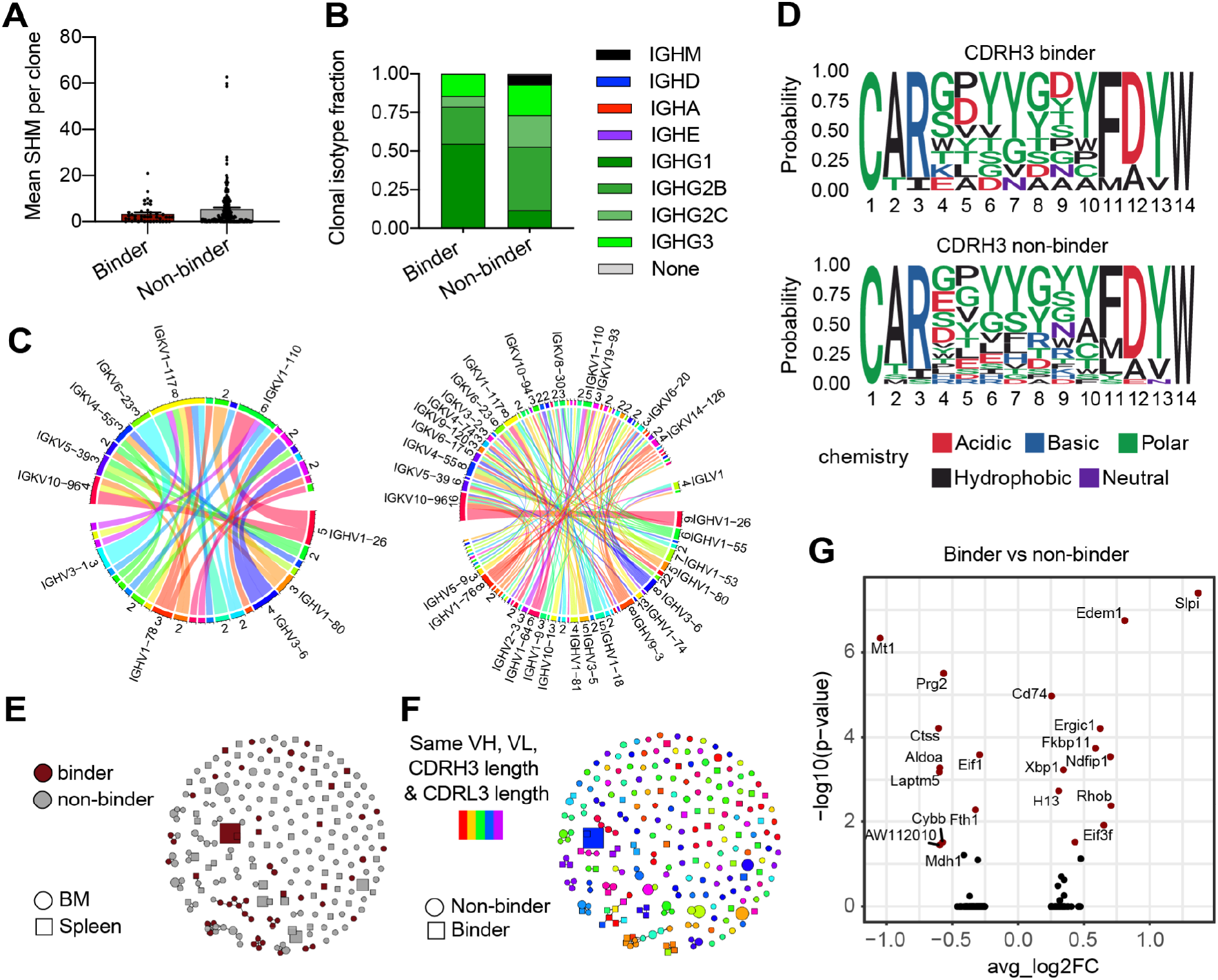
Investigating immune repertoire and transcriptional features of TNFR2-specific BM PCs. A. Mean number of nucleotide somatic hypermutations (SHMs) per TNFR2-specific and non-specific bone marrow clones. Mutations were calculated in the full-length V and J regions across both heavy (HC) and light chain (LC). Clone was determined by grouping those B cells containing identical CDRH3+CDRL3 amino acid sequences. B. Isotype distribution for the confirmed bone marrow TNFR2-specific and non-specific sequences on the clonal level. C, Sequence logo plots of the confirmed TNFR2-specific and non-specific CDRH3 sequences for the most frequent CDRH3 length. D. Circos plots depicting the relationship between HC and LC V genes for TNFR2 specific (left) and non-specific (right) sequences in the bone marrow. Color corresponds to the different V genes. Edges illustrate the number of clones using each particular combination. Only the most frequent V genes are noted. E. Similarity network of TNFR2-specific and non-specific B cell clones. Nodes represent unique clones. Edges connect those clones separated by an edit distance of 5 amino acids or less. Color corresponds to TNFR2 specificity. Shape indicates organ of origin. F. Same similarity network as (E) with color indicating clones with identical germline genes and CDR3 lengths. Shape indicates TNFR2 specificity. G. Differential gene expression between TNFR2 specific and non-specific clones in the bone marrow. Points in red indicate significantly differentially expressed genes (p-adj < 0.01).

We next investigated whether distinct gene expression profiles could be detected between TNFR2-specific and non-specific clones, as these signatures could correspond to genes involved in recent recruitment and migration compared to long-lived BM PCs present prior to immunization. Following differential gene expression analysis, we observed significantly expressed markers, both when including all TNFR2-specific sequences (Figure 4G) and when restricting our analysis to the different age cohorts (Figures S14A, S14B). This, however, did not reveal any meaningful differences in transcriptional signatures between antigen-specific and non-specific cells.

## Discussion

Here, we used single-cell antibody repertoire and transcriptome sequencing to investigate the extent that repertoire features can be leveraged to discover antigen-specific antibodies from immunized mice. scSeq allowed us to relate individual transcriptomes to the antibody repertoire for nearly one hundred thousand B cells, thereby providing large-scale insight into the relationship between gene expression, clonal selection, and antigen specificity. We observed comparable levels of clonal expansion in repertoires from both young and aged mice following serial immunizations with human TNFR2. In addition, we observed class-switched antibodies among the most expanded clones in both spleen and bone marrow repertoires.

The emergence of scSeq workflows has made it possible to comprehensively test and reconstruct the specificity of antibodies based on their immune repertoire profiles. Using antibody expression and screening, we were thus able to demonstrate that a fraction of expanded cells in the repertoires of young mice produced antibodies with specificity to TNFR2 following immunization. This fraction was comparable to our previous results in the context of immunization with OVA, where approximately 45% of the most expanded clones were found to be antigen-specific (Neumeier, Yermanos, et al. 2021). Moreover, our results demonstrate an increased proportion of clonally expanded B cells that were not antigen-specific in aged mice, suggesting that immunization-induced clonal expansion fails to exceed levels present in naive repertoires. This decrease of antigen specificity in aged mice may be linked to previous reports that naive B cell repertoires have restricted clonal diversity in aged individuals or that IgM+ B cells accumulate in the bone marrow during aging, thereby reducing available space for newly recruited B cells (Gibson et al. 2009; Miller and Allman 2003; Mehr and Melamed 2011). Other parameters linking immune senescence to a decreased number of antigen-specific plasma cells could involve alterations of pro-B/pre-B proliferative capacities (Min, Montecino-Rodriguez, and Dorshkind 2006; Shahaf, Johnson, and Mehr 2006), a decline of number and size of germinal centers (Zheng et al.1997) and deficiencies in class-switching (Frasca et al. 2004). Further measuring of affinities and, in the case of antiviral immunizations, neutralization potential, would be beneficial to assess how aging impacts the quality of the antibody response. In addition, we observed that antigen specificity was largely stochastic and could not be predicted based on the clonal rank of expanded cells in both young and old mice, which is in accordance with previous results (Neumeier, Yermanos, et al. 2021).

Having discovered TNFR2-specific sequences using clonal expansion and clonal overlap, we integrated single-cell transcriptomes with antibody repertoires in an attempt to investigate whether other factors could be incorporated into the selection criteria such as Ig isotype, SHM or transcriptional signatures. We focused on the BM PC repertoires due to the higher number of experimentally validated antibodies with specificity to TNFR2. These repertoires remain particularly interesting since plasmablasts, short- and long-lived PCs significantly contribute to circulating antibodies present in serum (Nutt et al. 2015). This comparison revealed that TNFR2-specific PCs had minor differences in immune repertoire features and similar transcriptional profiles to non-specific PCs. More specifically, our data suggested that antigen-specific clones were less mutated in the V- and J-regions and preferentially expressed the IgG1 subtype. We expected that immunization would lead to clear transcriptional differences relating to recent selection between TNFR2-specific and non-specific PCs. However, we observed minor transcriptional differences between these two populations of cells when performing unsupervised clustering and differential gene expression analysis. Transcriptional differences were however observed in IgM-, IgD-, IgA- and IgG-expressing B cells, which was consistent with previous results (Neumeier,Pedrioli, et al. 2021), as well as expansion-specific transcriptional clustering (Kuhn et al. 2021; Yermanos, Neumeier, et al. 2021).

Single-cell antibody repertoire sequencing provides a detailed molecular quantification of clonal selection. Thus far, the majority of repertoire studies conducted have examined lymphocytes following infection or immunization using model proteins (Bailey et al. 2017; Turner et al. 2020; Goldstein et al.2019; Burton and Hangartner 2016; Cao et al. 2020; Wen et al. 2020; Kräutler et al. 2020). In contrast, there is limited information regarding the behavior of humoral immunity triggered by an antigen with high host-sequence similarity, which are common therapeutic targets for indications such as cancer and autoimmune disease. Therefore our results have implications for the discovery of monoclonal antibodies targeting such therapeutic antigens, such as TNFR2, which is implicated in pro- and anti-inflammatory conditions (Medler and Wajant 2019; Fischer, Kontermann, and Pfizenmaier 2020).

**Table S1.Information on the sequences that were selected for antibody expression and specificity validation.** Antibody origin (mouse id), CDRH3+CDRL3 amino acid sequence (CDR3 aa), variable heavy and variable light nucleotide sequence (VH nt and VL nt respectively), reason for selection (selection criteria) and ELISA signal above background (ELISA fold-change) are shown. In the “selection criteria” column, overlapping sequences that were also expanded and used in figure 3 are labeled with a star (*).

## Methods

### Mouse experiments

All animal experiments were performed in accordance with institutional guidelines and Swiss federal regulations. Experiments were approved by the veterinary office of the canton of Basel-Stadt (animal experimentation permission 2582). 3-, 12- and 18-month-old C57BL/6 male (Janvier) mice were repeatedly immunized every 14 days (5 times) subcutaneously (s.c.) into the flank with 10μg of human TNFR2 protein (Peprotech, 310-12) and 20μg MPLA (Sigma, L6895) adjuvant in 150μL PBS.

### Isolation of bone marrow plasma cells and splenic B cells

A single cell suspension was prepared by flushing the bone marrow from the hind legs in RPMI containing 10% FCS buffer with 10 ng/mL IL6 (Peprotech, 216-16). A red blood cell lysis step was performed in 2 mL ammonium-chloride-potassium (ACK) lysis buffer for 1 minute at room temperature and subsequently inactivated with 20 mL RPMI containing 10% FCS. Single-cell suspensions were stained with the following antibodies in FACS buffer (1:200 dilution) CD138-APC, CD4-APC-Cy7, CD8a-APC-Cy7, NK1.1-APCCy7, Ter119-APC-Cy7, TACI-PE, B220-APC, CD19-PE-Cy7 for 30 minutes at 4°C. Cell sorting was performed using a FACSAria with FACSDiva software into RPMI.

### Single-cell sequencing of antibody repertoires

Single-cell sequencing libraries were constructed from the isolated BM PCs following the demonstrated 10x Genomics’ protocol: ‘Direct target enrichment - Chromium Single Cell V(D)J Reagent Kits’ (CG000166). Briefly, single cells were co-encapsulated with gel beads (10x Genomics, 1000006) in droplets using 5 lanes of one Chromium Single Cell A Chip (10x Genomics, 1000009) with a target loading of 13,000 cells per reaction. V(D)J library construction was carried out using the Chromium Single Cell 5’ Library Kit (10x Genomics, 1000006) and the Chromium Single Cell V(D)J Enrichment Kit, Mouse B Cell (10x Genomics, 1000072) according to the manufacturer’s instructions. Final libraries were pooled and sequenced on the Illumina NextSeq 500 platform (mid output, 300 cycles, paired-end reads) using an input concentration of 1.8 pM with 5% PhiX.

### Repertoire analysis

Raw sequencing files arising from Illumina sequencing lanes were supplied as input to the command line program cellranger (v3.1.0) on a high-performance cluster. Raw reads were aligned to the germline segments from the GRCm38 reference (vdj_GRCm38_alts_ensembl-3.1.0) and subsequently assigned into clonal families based on identical combinations of CDRH3+CDRL3 nucleotide sequence via cellranger. Further filtering was performed to include only those cells containing exactly one heavy chain and one light chain sequence. Firstly, clonotypes containing identical CDR3 amino acid sequences were merged into the same clonal family. Clonal frequency was determined by counting the number of distinct cell barcodes for each unique CDR3. Those cells in clones supported by only one cell were considered unexpanded clones, whereas those clones supported by two or more cells were considered expanded. Clone variants were determined as those sequences with the exact same VH and VL. SHMs were determined as nucleotide substitutions in the V and J regions. Clonal overlap was calculated based on identical amino acid CDRH3+CDRL3 sequences between any two or more repertoires. Germline gene usage was determined by cellranger’s vdj alignment to the murine reference. V and J gene combinations were calculated and visualized with circos plots using the VDJ_circos function in Platypus (v2.1) (Yermanos, Agrafiotis, et al. 2021) with a label.threshold of 2 or 5. Isotype was determined based on the constant region alignment per cell or for the majority of cells within each clonal family. In the case that the variable region alignment was provided but the isotype was not recovered, the isotype was labeled as “Unknown”. Jaccard indices were calculated by quantifying the intersection between two groups divided by the length of the union of the same groups. Similarity networks were calculated based on the VDJ_network function in Platypus (v2.1), which first calculates the edit distance separately for HC and LC CDR3s, and then draws edges between those clones with a distance below the specified threshold. ROC curves, AUC scores and confusion matrices were generated using the PlatypusML_feature_extraction_VDJ function in Platypus using the aa sequences of the CDRH3+CDRL3 regions. Feature importance was calculated using the xgb.ggplot.importance function from xgboost (v1.6.0.1).

### Transcriptome analysis

The output matrices, barcodes, and features files from cellranger (10x Genomics) for each sample were supplied as input to Seurat (v4.0) (Satija et al. 2015) using the Read10X function and subsequently converted into a Seurat object using the function “CreateSeuratObject”. Only those cells containing less than 20% of mitochondrial reads were retained in the analysis. All BCR related genes (V(D)J genes, isotype constant regions, J-chain) were filtered out prior to further analyses. Data was normalized using a scaling factor of 10000 and variable features were found using 2000 genes. Cluster resolution was set to 0.5 and the first fifteen PCR dimensions (determined via visualization of the normalized Seurat object using an elbow plot) were used for neighborhood detection. UMAP was performed again using the first fifteen PCA dimensions. All repertoires were proceeded and filtered together. Genes defining clusters were determined by setting the min.pct argument equal to 0.25 using Seurat’s “FindMarkers” function. For analyses involving B cells found in repertoire sequencing data, only those cells containing identical nucleotide barcodes sequences were included and reanalyzed in the Seurat object. Heatmaps of cluster defining genes were selected for the top genes ranked by logFC after employing the Bonferroni correction for multiple hypothesis testing (p.adj < 0.01).

### Antibody expression and validation

Antibodies were either transiently expressed in HEK 293 Expi cells using the ExpiFectamine 293 Transfection Kit (Thermo, A14524) and the pFUSE2ss vector system (Invitrogen) as previously described (Vazquez-Lombardi et al. 2018) or as stable hybridoma cell lines using CRISPR/Cas9 genome editing as before (Parola et al., 2019). Specificity was validated by performing normalized supernatant ELISAs against the extracellular domain of human TNFR2 (Peprotech, 310-12 and in-house production). Flow cytometry phenotyping of hybridoma cells was performed on a BD FACS Aria III. Typically, 5×10 cells were stained for 30 minutes on ice in 50 μl of a labeling mix consisting of anti-IgG2c-AlexaFluor488 (Jackson ImmunoResearch, 115-545-208) and anti-IgK-Brilliant Violet421 (BioLegend, cat# 409511) at 1:100 and 1:50 dilutions, respectively. Before acquiring, cells were washed twice. For ELISA experiments, 0.2 μm sterile-filtered cell culture supernatant of a 6d culture was used to confirm TNFR2 specificity. ELISA plates were coated with the capturing reagent (TNFR2 or control) in PBS at 4 μg/ml, blocked with PBS supplemented with 2% (w/v) milk (AppliChem, A0830) and incubated with cell culture supernatant. The supernatant from an influenza- (PDB 5GJT) and OVA-specific hybridoma served as negative controls (Neumeier, Yermanos, et al. 2021). A commercial antibody against human TNFR2 was used as a positive control (Invitrogen, AHR3022). An anti-mouse IgG-HRP (Sigma, A2554) was employed at 1:1500 and used for detection. Binding was quantified using the

1-Step Ultra TMB-ELISA substrate solution (Thermo, 34028) and 1M H_2_SO_4_ for reaction termination. Absorbance at 450 nm was recorded on an Infinite 200 PRO (Tecan). All commercial antibodies were used according to manufacturer’s recommendations. For each clone tested, the antibody variant (combined VH+VL nucleotide sequence) supported by the most unique cell barcodes was selected to be expressed. Clones were considered to be TNFR2-specific when the ELISA signal measured was at least three times higher than that of a negative background control unless otherwise specified.

### Hybridoma cell culture

Hybridoma cell lines were cultivated in high-glucose DMEM Medium (Thermo, 61965-026), supplemented with 10% (v/v) of ultra-low IgG FBS (Thermo, 16250078), 100 U/ml Pen/Strep (Thermo, 15140-122), 10 mM HEPES (Thermo, 15630-056) and 50 μM 2-mercaptoethanol (Thermo, 31350-010). Cell lines were maintained at 37 °C, 5% CO2 and passaged every 72 hours.

### Data visualization

Heatmaps depicting clonal overlap were created using the Rpackage pheatmap (v1.0.12). Heatmaps displaying differential gene expression were produced using the DoHeatmap function in the R package Seurat (v4.0) (Butler et al., 2018). Volcano plots and gene enrichment plots were ggplot (v3.3.5) (Wickham and Wickham, 2007). Similarity networks were produced using the R package igraph (v1.2.6) (Csardi et al., 2006). Sequence logo plots were generated using the R package ggseqlogo2 (v0.1) (Wagih, 2017). Experimental overview (Figure 1A) was created using BioRender.com. All other plots were produced using Prism v9 (Graphpad). All error bars indicate the standard error of mean.

## Acknowledgements

We acknowledge and thank Dr. Christian Beisel, Elodie Burcklen, and Ina Nissen at the ETH Zurich D-BSSE Genomics Facility Basel for support and assistance. We also thank the D-BSSE FACS facility for experimental support. **Funding**: This work was supported by the European Research Council Starting Grant 679403 (to STR), ETH Zurich Research Grants (to STR and AO), and an ETH Seed Grant (AY). **Competing Interests**: There are no competing interests.

---

**Figure S1.**
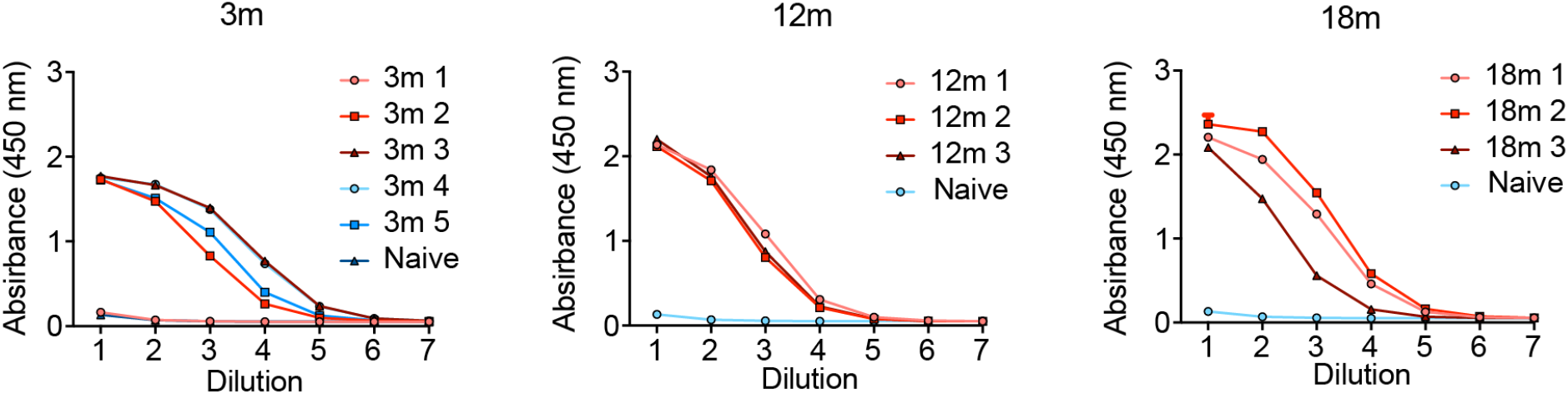
Serum antibody titers against hTNFR2 for each age group. Each line corresponds to a mouse. A naive mouse was included as a control.

**Figure S2.**
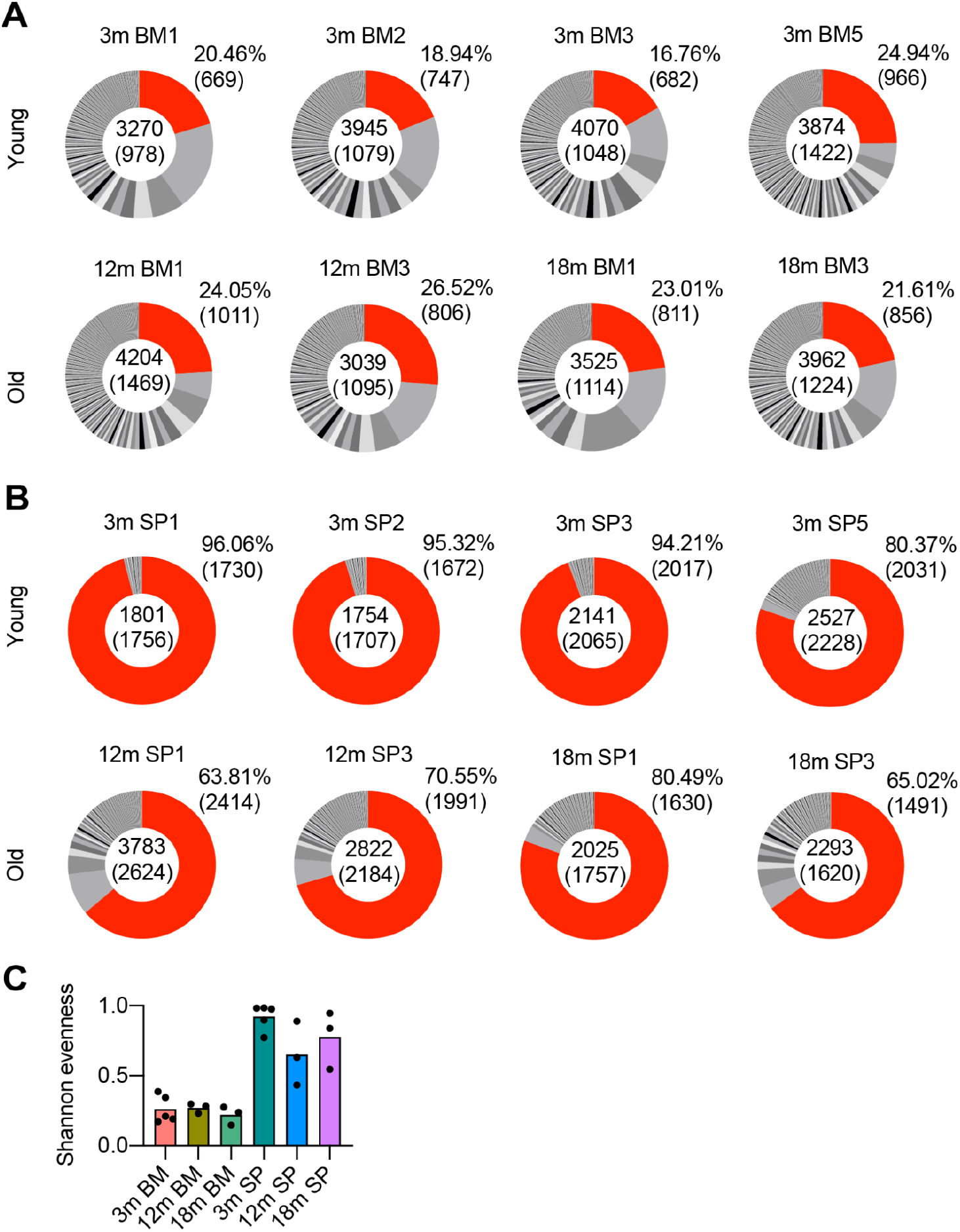
Clonal expansion following TNFR2 immunization. Distribution of clonal expansion in the A. bone marrow (BM) and B. spleen (SP). Each section corresponds to a unique clone and the size corresponds to the fraction of cells relative to the total repertoire. Red color highlights the fraction of clones containing 1 cell. Numbers in the center indicate the total number of cells and clones (in parenthesis). Numbers on the right indicate the percentage and total number (in parenthesis) of unexpanded clones. C. Shannon evenness quantifying clonal expansion of BM PCs and splenic B cells. Each individual point corresponds to a repertoire arising from a 3-month-old (3m), 12-months-old (12m), or 18-month-old (18m) mouse. Colors correspond to the different age and organ cohorts.

**Figure S3.**
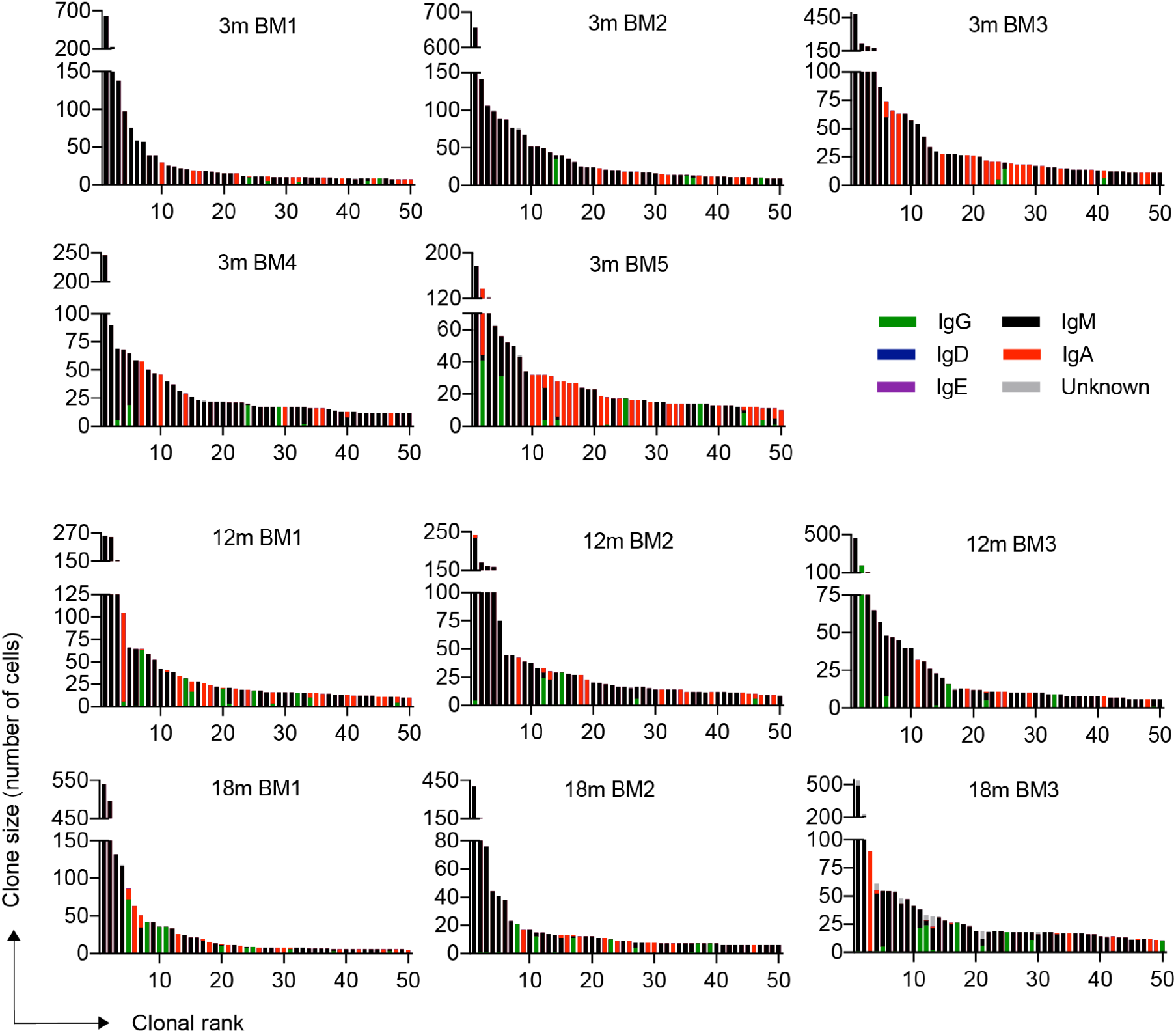
Clonal expansion for the top 50 most expanded clones of the BM PCs for each immunized mouse. Clones were determined by grouping those B cells containing identical CDRH3+CDRL3 amino acid sequences. Color corresponds to isotype. Each plot corresponds to a repertoire arising from a 3-month-old (3m), 12-months-old (12m), or 18-month-old (18m) mouse.

**Figure S4.**
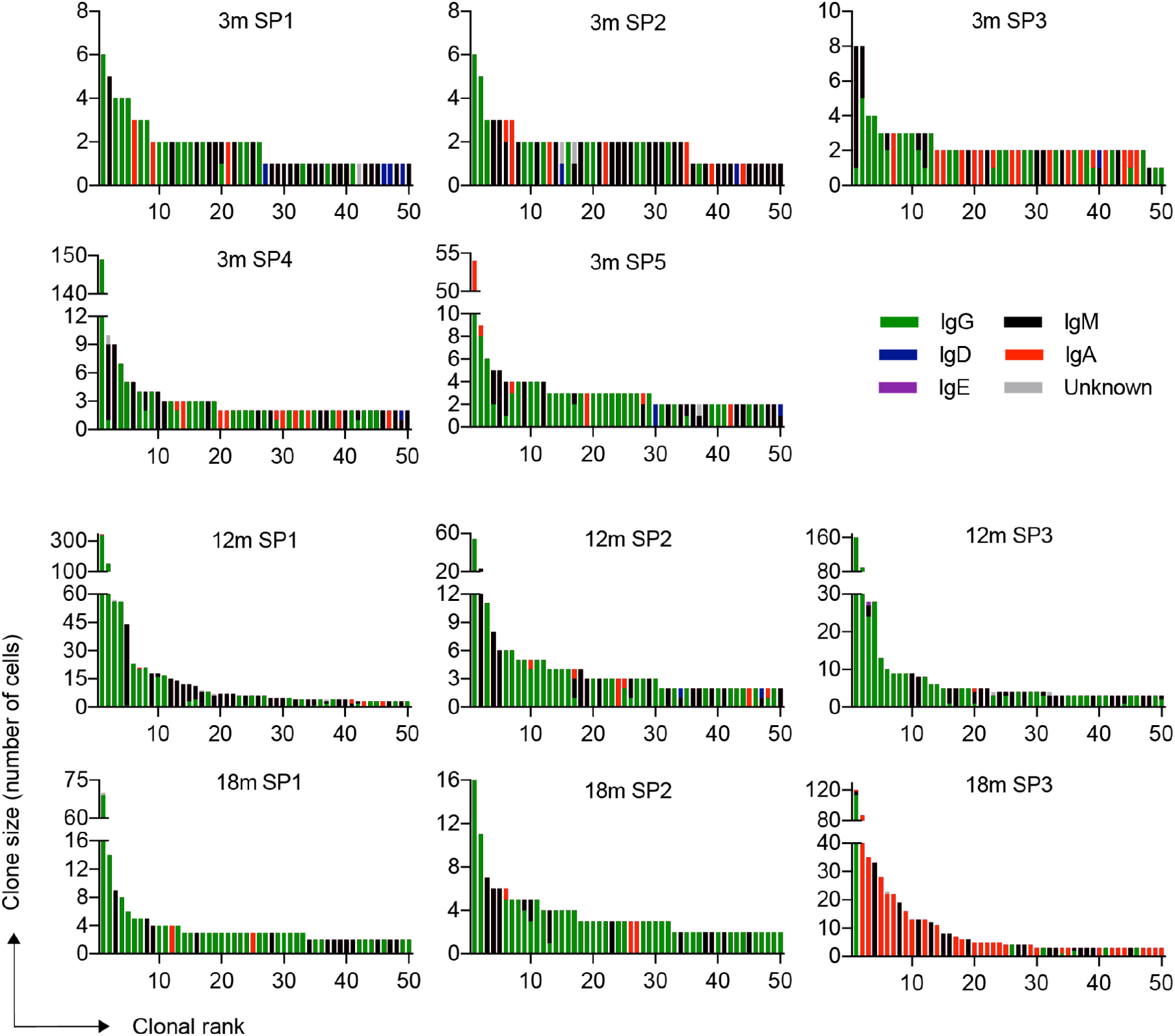
Clonal expansion for the top 50 most expanded clones of the splenic (SP) B cells for each immunized mouse. Clones were determined by grouping those B cells containing identical CDRH3+CDRL3 amino acid sequences. Color corresponds to isotype. Each plot corresponds to a repertoire arising from a 3-month-old (3m), 12-months-old (12m), or 18-month-old (18m) mouse.

**Figure S5.**
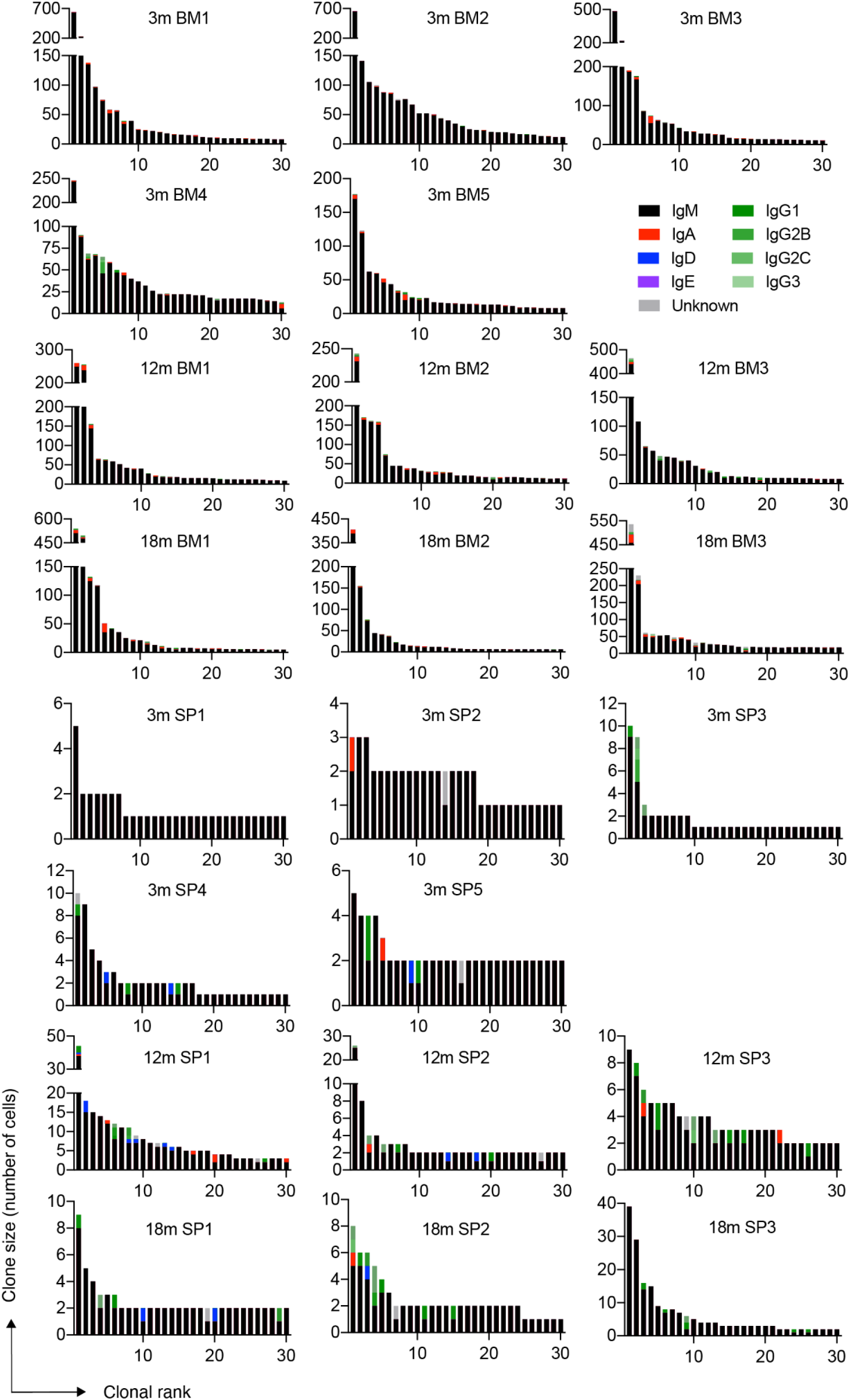
Clonal expansion for the top 30 most expanded clones with the majority of cells belonging to the IgM isotype of the BM PCs and splenic B cells for each immunized mouse. Clones were determined by grouping those B cells containing identical CDRH3+CDRL3 amino acid sequences.

**Figure S6.**
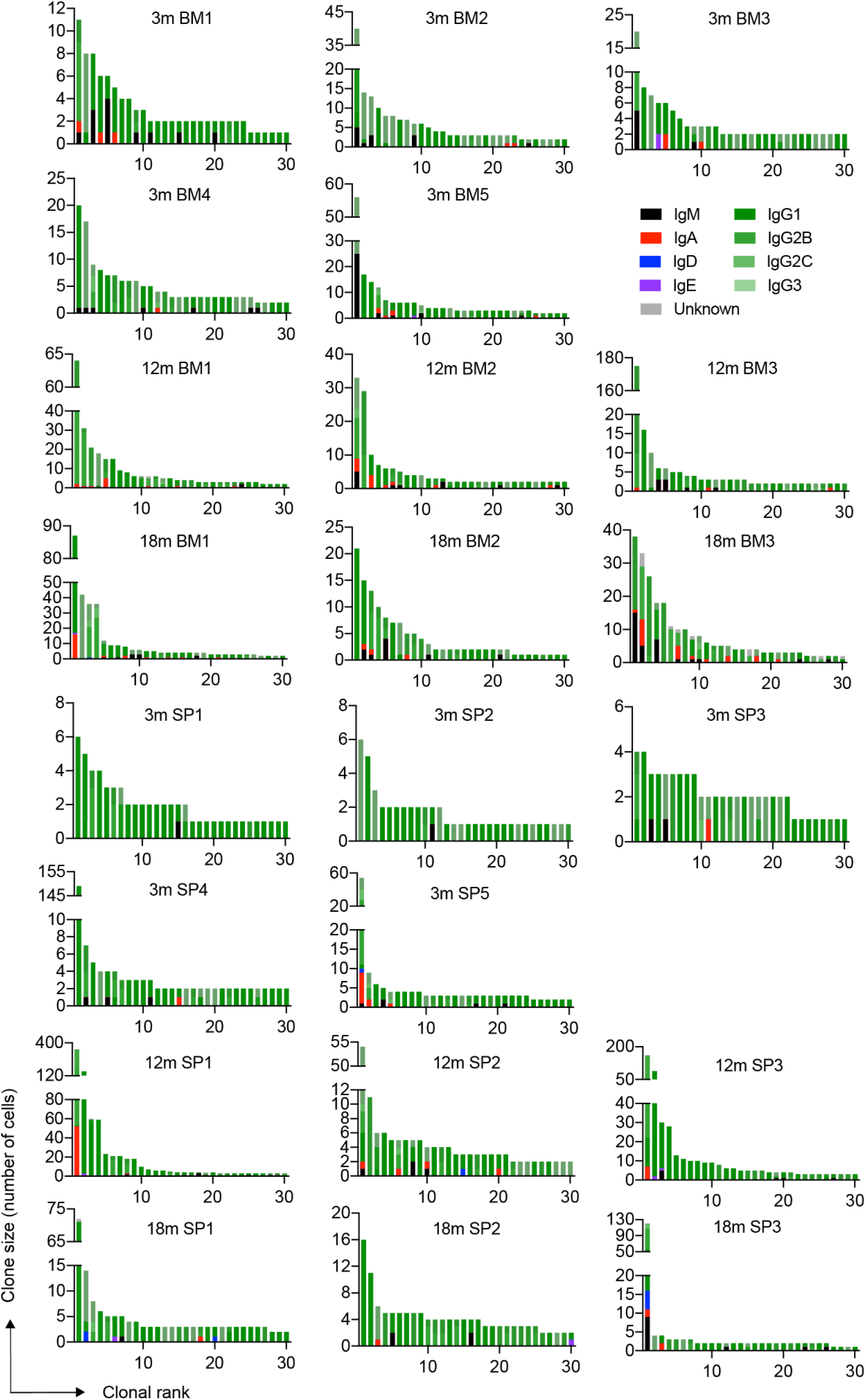
Clonal expansion for the top 30 most expanded clones with the majority of cells belonging to the IgG isotype of the BM plasma and splenic B cells for each immunized mouse. Clones were determined by grouping those B cells containing identical CDRH3+CDRL3 amino acid sequences.

**Figure S7.**
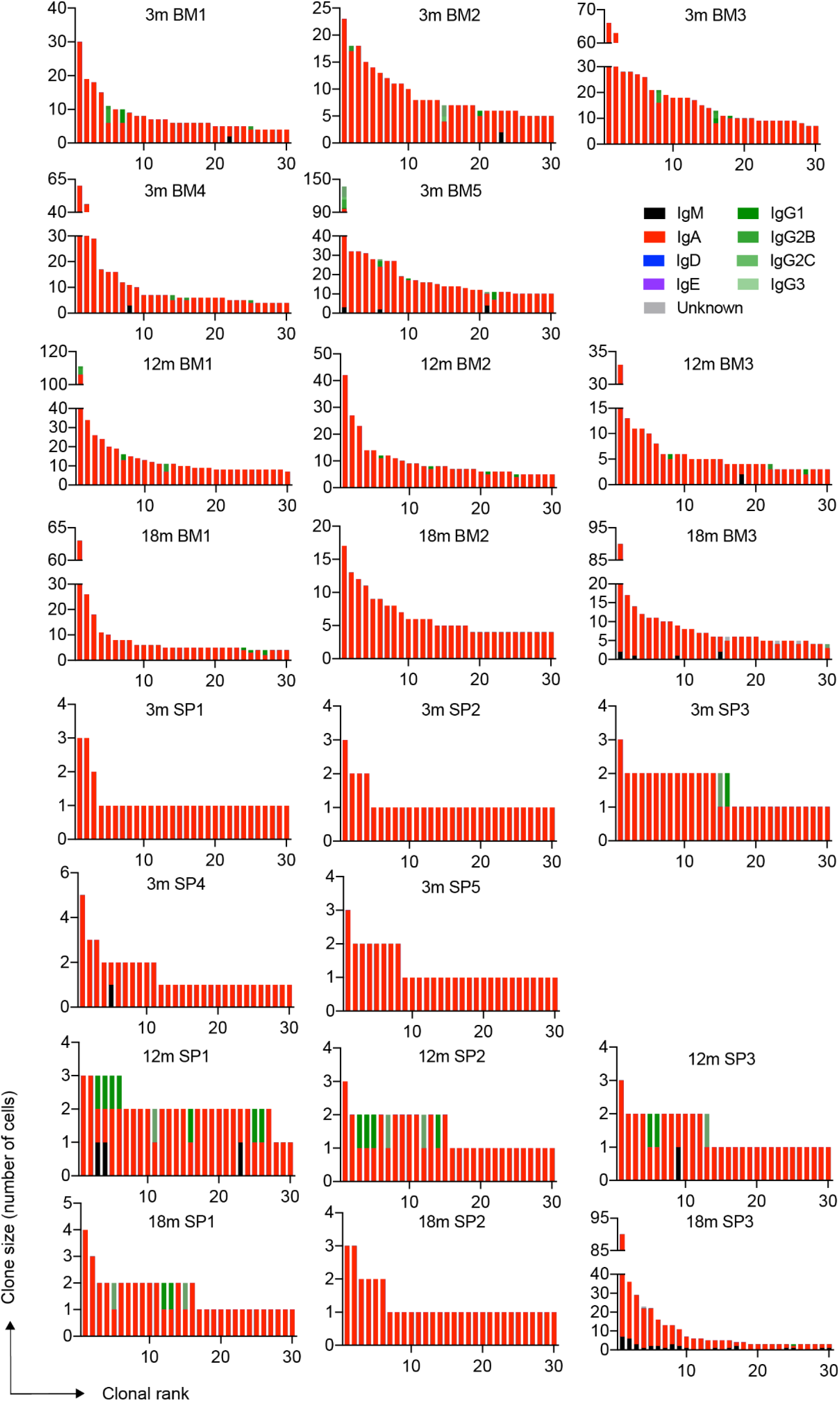
Clonal expansion for the top 30 most expanded clones with the majority of cells belonging to the IgA isotype of the bone marrow plasma and splenic B cells for each immunized mouse. Clones were determined by grouping those B cells containing identical CDRH3+CDRL3 amino acid sequences.

**Figure S9.**
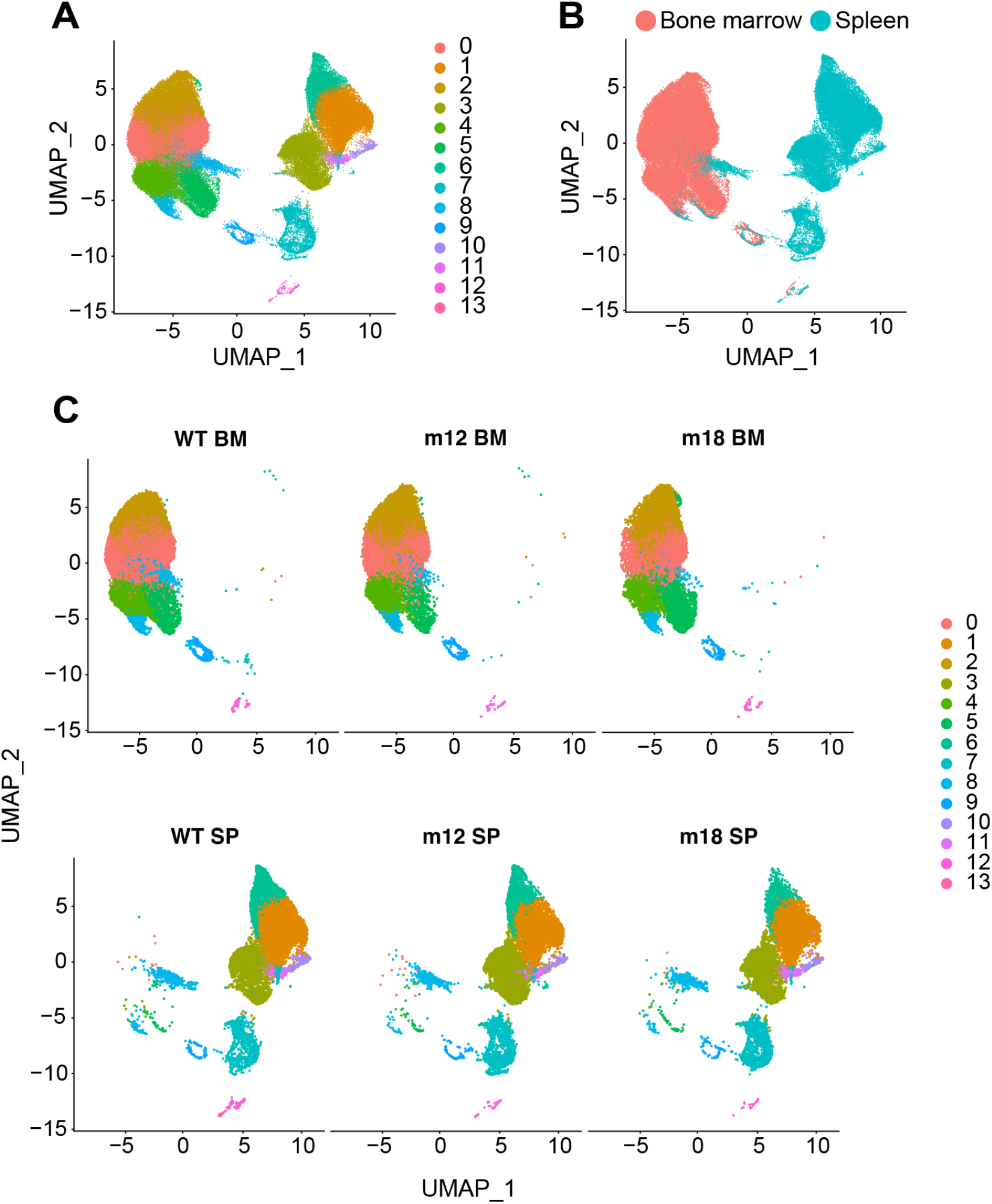
A. Uniform manifold approximation projection (UMAP) based on total gene expression of all repertoires following TNFR2 immunization. Each point corresponds to a cell and color corresponds to the transcriptional cluster. B. UMAP split by organ. C. Uniform manifold approximation projection (UMAP) split by age and organ cohort. Each point corresponds to a cell and color corresponds to the transcriptional cluster.

**Figure S10.**
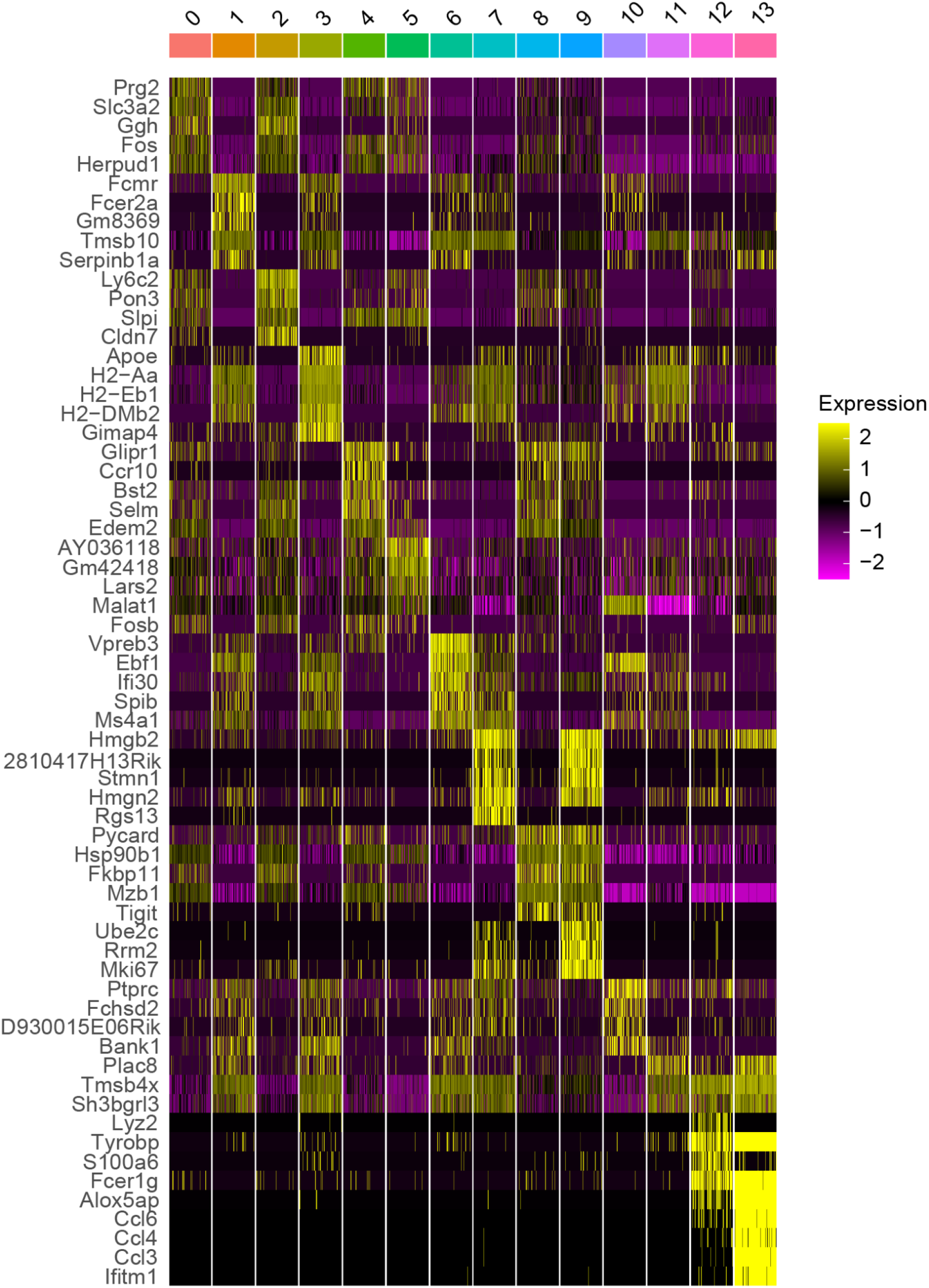
Top differentially expressed genes for each transcriptional cluster for all repertoires. Heatmap intensity corresponds to normalized expression. Each column represents a single cell and each row corresponds to a single gene. The top five genes based on average log fold change (avg_log2FC) have been selected for each cluster after removing ribosomal protein L (RPL), ribosomal protein S (RPS), and mitochondrial genes. All displayed genes had an adjusted p value less than or equal to 0.01.

**Figure S11.**
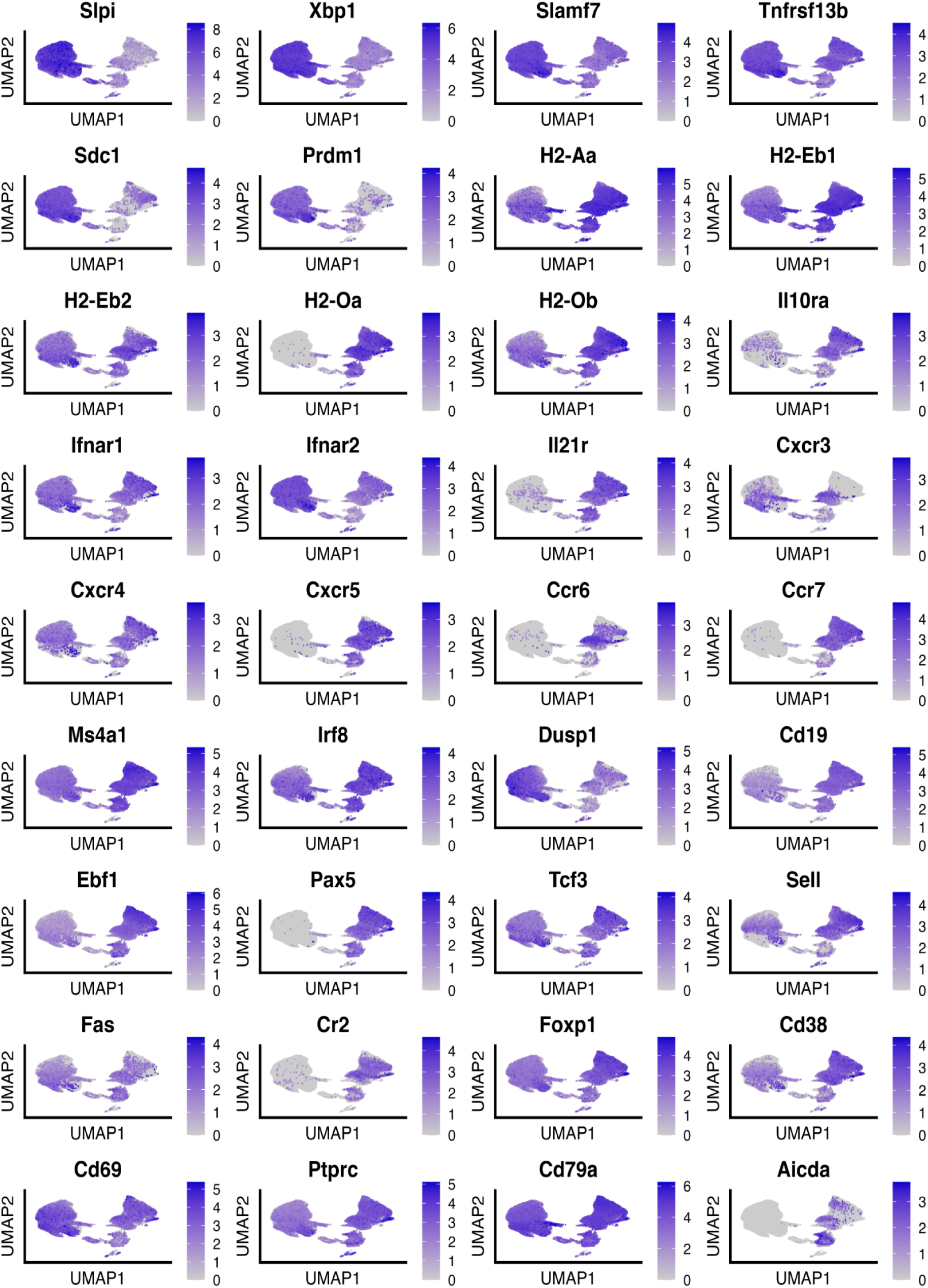
Uniform manifold approximation projection (UMAP) plots showing gene expression for selected genes.

**Figure S12.**
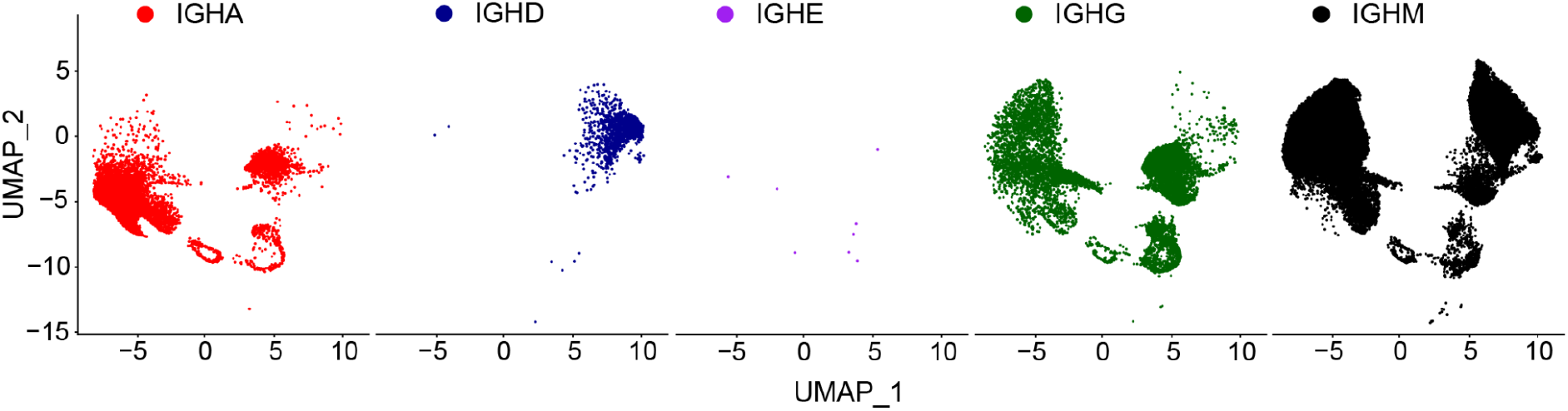
Uniform manifold approximation projection (UMAP) split by isotype. Each point corresponds to a cell and color corresponds to the respective isotype.

**Figure S13.**
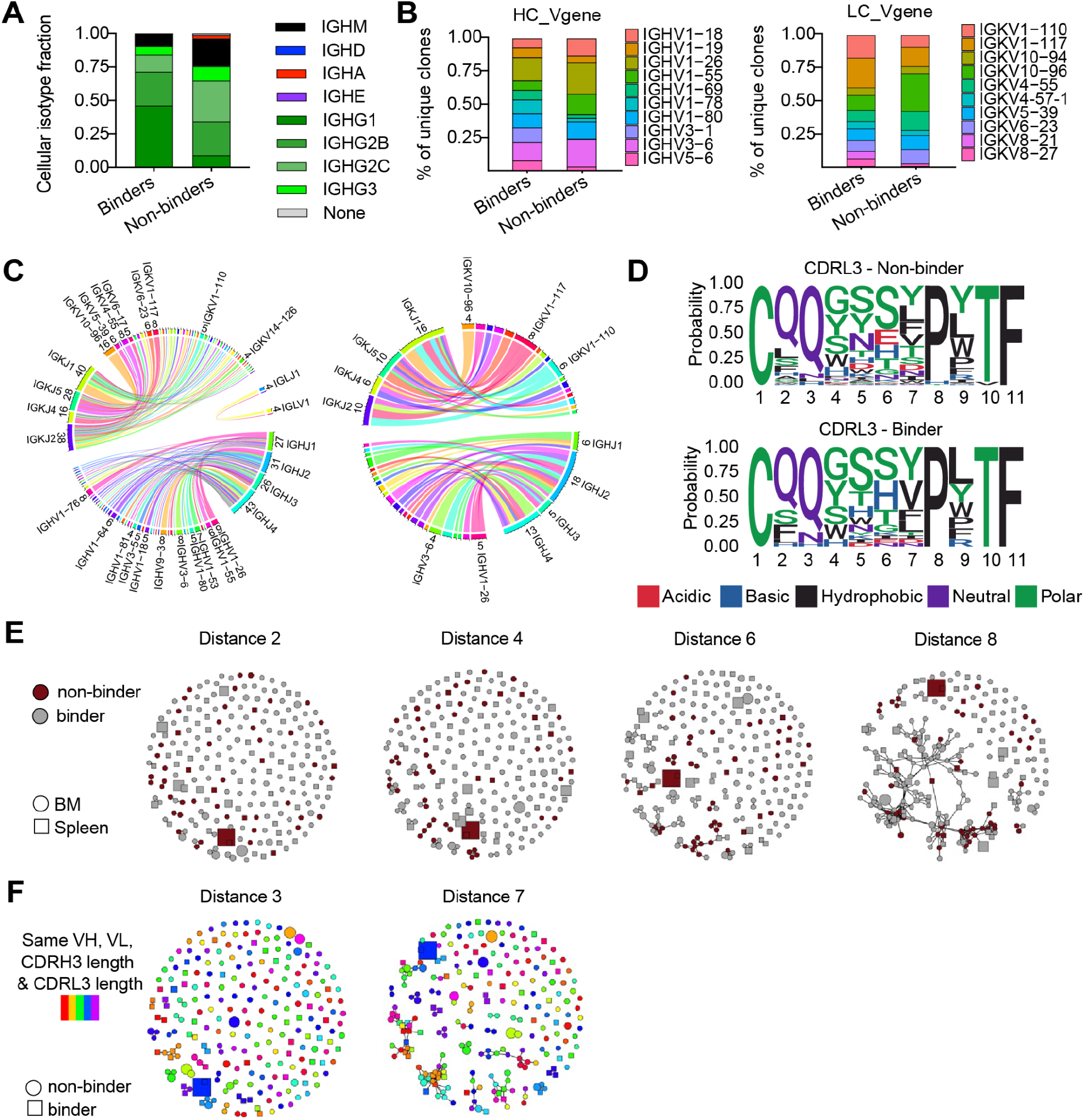
Immune repertoire and transcriptional features of TNFR2-specific sequences. A. Isotype distribution for the confirmed bone marrow TNFR2-specific and non-specific sequences on the cellular level. B. Sequence logo plots of the confirmed TNFR2-specific and non-specific CDRL3 sequences for the most frequent CDRL3 length. C. Heavy chain (right) and light chain (left) V gene distribution between antigen-specific and non-specific clones. D. Circos plots depicting the relationship between HC and LC V and J genes for TNFR2 specific (left) and non-specific (right) sequences in the BM. Color corresponds to the different V genes. Edges illustrate the number of clones using each particular combination. Only the most frequent V genes are noted. E. Similarity network of TNFR2-specific and non-specific clones. Nodes represent a unique clone. Edges connect those clones separated by an edit distance of 2, 4, 6 and 8 amino acids or less. Color corresponds to TNFR2 specificity and shape indicates organ of origin. F. Same similarity network as (E) with color indicating same variable HC, variable LC, CDRH3 length and CDRL3 length. Shape indicates TNFR2 specificity.

**Figure S14.**
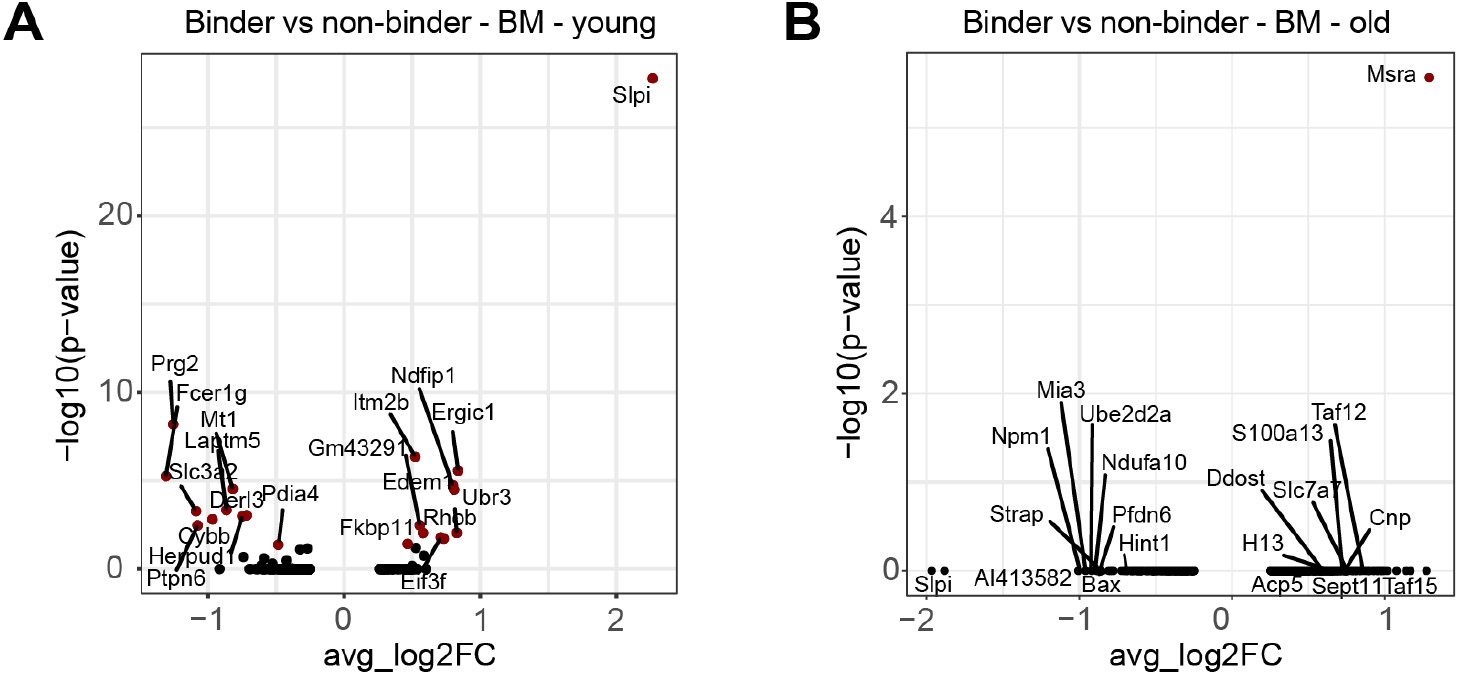
Differential gene expression between TNFR2 specific and non-specific clones in the bone marrow of young (A) and old (B) mice. Points in red indicate significantly differentially expressed genes (p-adj < 0.01).

